# The Dynamic Shift Detector: an algorithm to identify changes in parameter values governing populations

**DOI:** 10.1101/522631

**Authors:** Christie A. Bahlai, Elise F. Zipkin

## Abstract

Environmental factors interact with internal rules of population regulation, sometimes perturbing systems to alternate dynamics though changes in parameter values. Yet, pinpointing when such changes occur in naturally fluctuating populations is difficult. An algorithmic approach that can identify the timing and magnitude of parameter shifts would facilitate understanding of abrupt ecological transitions with potential to inform conservation and management of species.

The “Dynamic Shift Detector” is an algorithm to identify changes in parameter values governing temporal fluctuations in populations with nonlinear dynamics. The algorithm examines population time series data for the presence, location, and magnitude of parameter shifts. It uses an iterative approach to fitting subsets of time series data, then ranks the fit of break point combinations using model selection, assigning a relative weight to each break. We examined the performance of the Dynamic Shift Detector with simulations and two case studies. Under low environmental/sampling noise, the break point sets selected by the Dynamic Shift Detector contained the true simulated breaks with 70-100% accuracy. The weighting tool generally assigned breaks intentionally placed in simulated data (i.e., true breaks) with weights averaging >0.8 and those due to sampling error (i.e., erroneous breaks) with weights averaging <0.2. In our case study examining an invasion process, the algorithm identified shifts in population cycling associated with variations in resource availability. The shifts identified for the conservation case study highlight a decline process that generally coincided with changing management practices affecting the availability of hostplant resources.

When interpreted in the context of species biology, the Dynamic Shift Detector algorithm can aid management decisions and identify critical time periods related to species’ dynamics. In an era of rapid global change, such tools can provide key insights into the conditions under which population parameters, and their corresponding dynamics, can shift.

**Author Summary:** Populations naturally fluctuate in abundance, and the rules governing these fluctuations are a result of both internal (density dependent) and external (environmental) processes. For these reasons, pinpointing when changes in populations occur is difficult. In this study, we develop a novel break-point analysis tool for population time series data. Using a density dependent model to describe a population’s underlying dynamic process, our tool iterates through all possible break point combinations (i.e., abrupt changes in parameter values) and applies information-theoretic decision tools (i.e. Akaike’s Information Criterion corrected for small sample sizes) to determine best fits. Here, we develop the approach, simulate data under a variety of conditions to demonstrate its utility, and apply the tool to two case studies: an invasion of multicolored Asian ladybeetle and declining monarch butterflies. The Dynamic Shift Detector algorithm identified parameter changes that correspond to known environmental change events in both case studies.

## Introduction

Abrupt and persistent changes in ecological processes, and methods to detect them, have long interested ecologists [1–5]. Changes to the rules governing system dynamics are often associated with substantial changes to biodiversity and ecosystem functions. Understanding when, and how these changes occur is critically important to broader evaluation of system behaviors. The study of abrupt changes, discontinuities, and regime shifts is highly interdisciplinary, and has been examined for a variety of processes related to climate [6,7], ecology [8], and economic and social systems [9,10].

Population dynamics are determined by internal, biotic rules and also external abiotic factors, leading to both stochastic and deterministic forces affecting abundance through time [11]. External perturbations can lead to shifts in population dynamics, such that the parameters governing population abundance transition to other values [12,13]. In the context of this study, we define the set of parameters controlling dynamics as a population’s *dynamic rule*, and an abrupt shift in these parameter values as a *dynamic shift*. We use the term *break point* to describe the location in time series data where the dynamic shift occurs.

Although theoretically straightforward, identifying dynamic shifts in noisy ecological systems is challenging using real-world data due to a lack of systematic, adaptable tools [2]. Arguably, the most common approach to identify break points is through the use of segmented regressions [12,14,15]. However, these models don’t adequately account for nonlinearities, and uncertainties in the existence and location of breaks are typically not quantified [1,2]. Break points are often applied to time series data *ad hoc*, based on data visualization or specific hypotheses surrounding factors affecting population changes [12,14–17], creating the potential for biases in break point selection.

Several break point detection methods have been developed to address issues associated with *ad hoc* approaches. Such dynamic shift analysis tools use a variety of statistical optimization strategies, including linear and moving average methods [18–21]. For example, climatological and econometrics time series data are frequently examined for stepwise statistical deviations from the mean or variance [6,7,22]. To fit the periodicity of population time series data, wavelet analyses have also been used to detect break points [23] but this method does not mechanistically account for density-dependent processes such that changes in parameter values are not easily interpretable [24]. Dynamic shift detection methods that explicitly account for non-linear population processes may be less likely to yield false positives than methods based on summary statistics [25].

Break point detection methods based on statistical measures tend to rely on null hypothesis testing (i.e., that no dynamic shift occurred) and thus they have low sensitivity in situations where statistical power is limited. Additionally, such methods do not provide a means for assessing uncertainty in the existence and magnitude of break points [1,26]. In a 2009 review, Andersen and colleagues note that if common break point methods were used on typical ecological time series with 20-40 time steps, only the most extreme transitions occurring near the midpoint of the time series would be deemed ‘significant’ [1]. They concluded that break point analyses could be enhanced with respect to both sensitivity and parsimony by use of model selection procedures. Thus, to address these limitations in the ability to identify dynamic shifts in population processes, it is necessary to develop rigorous tools that allow users to accommodate non-linear population models and quantify uncertainties associated with the existence of potential break points.

In this paper, we develop a generalizable algorithm, the Dynamic Shift Detector (DSD), to identify dynamic shifts in populations with density-dependent growth using time series data. The DSD algorithm uses an iterative approach, grounded in information theoretic model selection. We illustrate the DSD using the Ricker model because of this model’s simplicity and high performance under a variety of realistic environmental scenarios. Density-dependent population models such as the Ricker provide a convenient, generalizable proxy for a variety of ecological processes because of their relatively simple parameterization and potential to explain complex dynamics [27]. Although deterministic approaches to population modelling have largely fallen out of favor for more complex structures and stochastic elements [28–30], simple dynamic models remain useful due to their easily interpretable and ecologically meaningful parameters [31]. Further, the techniques described in our paper can be readily adapted to other model structures, including more complicated processes such as seasonal periodicity or lag effects.

We describe the basic structure of our DSD algorithm and how it can be used to evaluate the presence, location, and magnitude of dynamic shifts in population parameters (i.e., break points in time series abundance data). We demonstrate the utility of our algorithm through a series of simulations and apply the algorithm to empirical case studies of two populations of economic and conservation concern. First, we examine the invasion process of the multicolored Asian ladybeetle (*Harmonia axyridis*), a cosmopolitan invasive, in the two decades following its arrival in Midwestern US agricultural ecosystems. Then, we examine the declining eastern monarch butterfly (*Danaus plexippus*) population using census data collected on its overwintering grounds in Mexico over a similar two-decade period. In our ladybeetle case study, the DSD algorithm identified dynamic shifts associated with known variation in prey availability, with moderately high weight for a break coinciding with prey arrival and moderately low weight for a break coinciding with management actions aimed to control the prey. The results for the monarch population were more ambiguous, with greater uncertainty about the number and location of breaks in the time series. Several equivalently performing break point combinations had divergent weights for various break points, suggesting that multiple, super-imposed biological processes drive the dynamics of this population.

## Models

### The Dynamic Shift Detector algorithm

Although any time series population model can be used with our tool, we illustrate the DSD algorithm with a Ricker population model. To do this, we assume that the population size in time t+1, *N*_*t*+1_, is dependent on the population size in time t, *N*_*t*_ and regulated by two parameters: the carrying capacity of the system, *K*, and the annual intrinsic growth rate, *r* [27]:

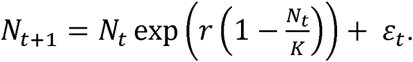

We further assume that observed annual population abundance is partially stochastic and may be influenced by environmental variation and sampling error. We include an error term *ε*_*t*_ to represent this noise, which follows a normal distribution centered around zero with a variance of *σ*^2^. The parameters K, r, and *ε*_*t*_ are estimated from the population time series data (N_1_, N_2_, … N_*t*_). The Ricker model is a useful starting point for break point analyses because 1) it does not rely on any external information (abundance in time t is a function of only abundance in time *t*-1); 2) only three parameters (including the error) need to be estimated, and those parameters have ecologically meaningful interpretations; and 3) it is an extremely flexible distribution, taking a variety of forms, from linear to compensatory to over-compensatory, and thus has a wide range of applications across a variety of taxa [32,33]. Subsequent applications of the DSD algorithm can incorporate other population models if the life history of the target organism requires a different structure.

To build the DSD algorithm, we use an iterative, model-selection process to determine if, and when, shifts in parameter values occur within a given time series. To achieve this, we first fit the Ricker model to the entire time series of available data. Then the population time series is subdivided into all possible combinations of 2, 3, …, n subsets of sequential data points (hereafter, ‘break point combinations’) and the Ricker model is fit to each of the subsets produced for each break point combination. To avoid overfitting, we constrain break point combinations to include only subsets with a minimum of four sequential data points.

After fitting all break point combinations, we evaluate the candidate set of models by calculating the Akaike Information Criteria for small sample sizes (AICc) for each segment and summing them accordingly [34]. Fits for break point combinations with comparatively lower AICc values are considered to have better performance. For a given time series, the DSD algorithm produces a set of top performing break point combinations for cases in which model fits produce equivalent AICc values (i.e. within 2 units of the best-performing fit; [35]). To evaluate the strength of evidence for an identified break in the time series, we use the relative variable importance method [35]. We compute the Akaike weight *w*_i_ (a measure of the relative likelihood of a break point combination, given the data and the set of break point combinations being tested) for every identified break point across all combinations. Commonly used in model averaging, the *w*_1_, *w*_2_,… *w*_n_ are interpreted as the respective conditional probabilities for each model in a set of *n* models [36]. Break weight (i.e., relative variable importance, *sensu* [35]) is computed as the sum of the Akaike weights for all break point combinations, where that break point appears. Break point combinations with weights <0.001 were excluded to increase computational efficiency.

We selected AICc as our information criterion for model selection within the DSD algorithm because it provides a balance of specificity and sensitivity. However, we also completed a parallel analysis with an identical procedure using AIC as the information criterion for decision-making, which is documented in Appendix S1. AICc is a function of AIC with a correction for small sample bias, which is appropriate for the sample sizes typical to contemporary population time series data (i.e., 15-30 years/data points) and is designed to minimize the risk of overfitting during model selection [35]. However, use of AIC for model selection may be desirable when increased algorithmic sensitivity to dynamic shifts is desired.

The DSD algorithm is implemented as a series of R functions to enable a user to quickly generate a list of potential break points for a population time series dataset. The algorithm (and all subsequent simulations and case studies) were scripted and run in R Version 3.3.3 [37]. For fitting the Ricker model, we used the Levenberg-Marquardt nonlinear least-squares algorithm as implemented in the package minpack.LM [38]. All data manipulations, analyses and figure scripts, including the complete development history, are publicly available in a Github repository at https://github.com/cbahlai/dynamic_shift_detector [39]. We summarize the role of each function used in the algorithm within Appendix S2.

### Simulation study

We conducted a series of simulations to test the accuracy of the DSD algorithm under a variety of plausible parameter spaces. For all scenarios, we fix *N*_*1*_ = 3000, and *K* = 2000 in the initial conditions, as the Ricker model is most reliably fit for populations fluctuating around their carrying capacity. As the dynamic observed in a Ricker population is driven primarily by the relationship of other parameters to *K* than by the absolute value of *K* itself, we held the starting value of *K* constant for all simulations. For each set of simulations, we held the variables not being varied at “base values” defined as: starting value of *r* = 2, change in *r* = ±25%, change in *K* = ±75%, 2% noise (*τ*; described below), with time series length of 20 years. We examined the effect of the size of initial *r* on algorithm performance by creating scenarios with different starting values of *r* = 0.5, 1, 1.5, 2. For each value of initial *r*, we modified the percent change in r at break points from the starting values (± no change, 10%, 25%, 50%, 75%) while holding all other parameters at base values. We then ran a set of simulations examining the percent change in K at break points from its starting value (± no change, 10%, 25%, 50%, 75%) while holding all other parameters (including *r*) at base values. This led to a total of 40 scenarios (four starting values of *r* with five percent changes in *r* and five percent changes in *K*).

We further evaluated how the magnitude of stochasticity in the system (as measured by the error term *ε*_*t*_) influenced algorithm performance. For generalizability of our simulation results, we simulated error as a percentage of the mean population size, rather than as absolute value (as described in the model above that we used for fitting the DSD). For each annual population size in the simulated dataset, a random value was selected from a normal curve of mean 0 and standard deviation of *τ* (where *τ* = 1%, 2%, 5%, 10%, 15%) and multiplied by the expected population size generated from the deterministic portion of the model.

We ran these simulations with all noise (*τ*) levels across all percent change values for *r* and *K* (with other parameters held at base values) for a total of 50 additional scenarios (five percent noise values with five percent changes in *r* and five percent changes in *K*). Finally, we tested the impact of time series length by modifying the length of the simulated time series at five-year intervals over a range from 15 – 30 years (as the number of break point allows) while holding all other parameters constant, for four additional scenarios. We generated 250 simulated datasets for each of the 94 possible scenarios assuming breakpoint combinations with 0, 1, 2 and 3 breaks. Break point locations were randomly selected from within the set of possible time points. In total, we generated 93,572 data sets that we examined with our DSD algorithm. (Note that 94,000 simulations were run but simulations for higher numbers of break points in shorter time series occasionally failed to converge; results for such combinations are not presented).

We evaluated the DSD algorithm’s performance for all test scenarios by examining its ability to identify the true break points within the set of the best fitting break point combinations (i.e. the top ranked break point combination and those break point combinations whose AICc values fell within two units of the top ranked). We also examined the performance of the break-point weighting tool by calculating the average weightings of all true and erroneous break points identified in the top performing model(s) across all runs of a given scenario.

## Results

### Simulation Study

The scenario with the correct number of breaks and their locations was detected within the top performing break point combination sets with >70% accuracy under nearly all parameterizations (Fig. 1). Accuracy was generally lowest in time series with three break points but above 70% for most scenarios. These results remained roughly consistent regardless of the value of the variance (σ^2^) determining the annual amount of environmental/sampling noise (Fig. 1 A). Results were similar across all r values tested but performance of the DSD declined slightly when initial r was large (>2.0; Fig. 1 B). The DSD algorithm had the highest accuracy with larger shifts in K (≥25%; Fig 1. C) and relatively smaller changes to *r* (≤25%; Fig. 1 D). This result is somewhat counter-intuitive, as we would generally expect large shifts in all parameters to be more easily detected. However, because nonlinear models produce chaotic dynamics with high population growth rates (e.g., r > 2.3 in the Ricker model), a large shift in parameters could potentially result in a situation where multiple break point fits would perform equally well. Finally, the accuracy of the DSD algorithm decreased as scenario length increased, likely because of the factorial increase in potential break point combinations with additional data in the time series (Fig. 1 E). Accuracy was also lower in cases where the number of break points was high relative to the time series length (e.g., 20 years and three breaks).

**Figure 1:**
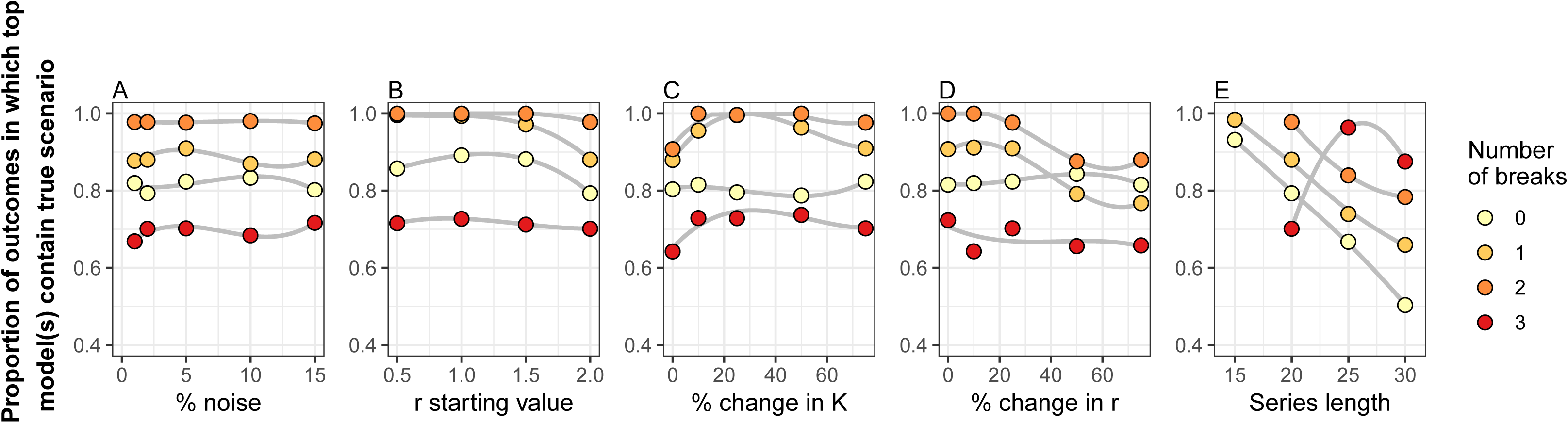
Performance of the Dynamic Shift Detector (DSD) algorithm under varying parameter values. Proportion of simulation results in which the true break scenario was detected within the top break point combinations as identified by the DSD implemented with an underlying Ricker model with varied A) noise (in the form of normally distributed error), B) starting values of the *r* parameter, C) percent changes in the *K* parameter, D) percent changes in *r*, and E) simulated time series length. Sets of 0, 1, 2 and 3 break points were randomly generated from within the set of possible values, and 250 datasets were simulated for each scenario. In each panel, other variables (that were not being varied) were held constant at their base values (i.e., noise=2%; starting value of *r* = 2; change in *r* = ±25%; change in *K* = ±75%; time series length = 20 years). Trends within a set of scenarios (grey lines) are illustrated with a third-order GAM smoothing line.

The breakpoint weighting analysis revealed that in the vast majority of cases, the average weight of a true break exceeded a value of 0.8 (Fig. 2A-E), whereas the weight of erroneous breaks averaged less than 0.2 in weight. The notable exception occurred when true breaks resulted from very small shifts in K (Fig. 2 C). Thus, our results suggest the following decision rules to evaluate strength of evidence for a break occurring at a given time point: when a weight of >0.8 is indicated for a break found by the DSD algorithm, we can reasonably conclude this is a true break, and likewise, a break with a weight of <0.2 can reasonably assumed to be erroneous. Weight values intermediate to those two thresholds can be interpreted as a quantification of the strength of evidence that a break occurred.

**Figure 2:**
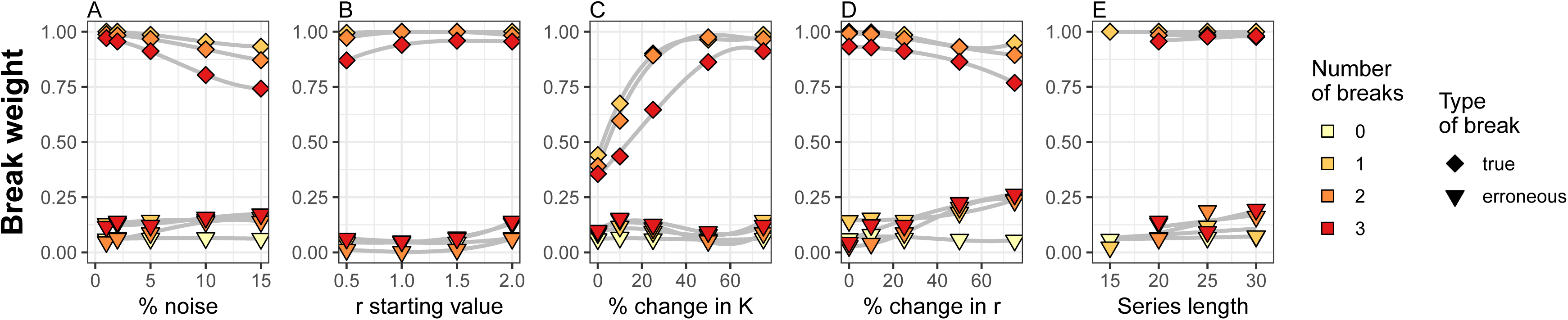
Average break weight of break points detected under varying parameterization conditions. Average weights of break points identified by the Dynamic Shift Detector algorithm reflecting true parameterization conditions (diamonds) or erroneous breaks suggested by the algorithm (triangles) under varied A) noise (in the form of normally distributed error), B) starting values of the *r* parameter, C) percent changes in the *K* parameter, D) percent changes in *r*, and E) simulated time series length. Sets of 0, 1, 2 and 3 break points were randomly generated from within the set of possible values, and 250 datasets were simulated for each scenario. In each panel, other variables (that were not being varied) were held constant at their base values (i.e., noise=2%; starting value of *r* = 2; change in *r* = ±25%; change in *K* = ±75%; time series length = 20 years). Trends within a set of scenarios (grey lines) are illustrated with a third-order GAM smoothing line.

### Case study applications

We tested the performance of the DSD algorithm with two case studies using population time series data from field observations. Both case studies involve approximately two decades of observations of economically or culturally important insect species: one examines an invasion process and the other examines a population decline, both occurring over the same time period in recent history.

#### Multicolored Asian ladybeetles in southwestern Michigan

The 1994 invasion of multicolored Asian ladybeetles to southwestern Michigan, United States was documented in monitoring data collected on agriculturally-important Coccinellidae (ladybeetles) in landscapes dominated by field crops. Population density of ladybeetles is monitored in ten plant communities weekly over the growing season using yellow sticky card glue traps starting in 1989 at the Kellogg Biological Station at Michigan State University. We used data on the captures of adults at the site from 1994-2017, culled at day of year 222 (August 10) to minimize the effect of year-to-year variation in the sampling period. We then calculated the average number of adults captured per trap, across all traps deployed within a sampling year, and used this value in our analysis. Detailed sampling methodology is available in previous work [40–42].

Two break points, one occurring after 2000 and one occurring after 2005, were observed in the top performing break point combination (Fig. 3 A, AICc=−18.02). However, the DSD algorithm indicated that two additional break point combinations, a single break after 2000 (AICc=−17.46), and a no break series (AICc=−17.64), had equivalent performance. Break weight analysis suggested a weight of 0.56 for the 2000 break, and a weight of 0.29 for the break after 2005. As these weights fall into a range intermediate to the 0.2 and 0.8 decision rules, we conclude that there is moderately strong evidence of a shift in dynamic rule after 2000, and moderate-weak evidence for a shift after 2005. The shift in 2000 is characterized by substantial increases in the values of *K* and *r*, with approximate increases of 75% and 40% over their initial estimates, respectively (Table 1). The shift in 2005 is characterized by a return to parameter estimates that were nearly identical to those observed at the beginning of the time series (Table 1, Fig. 3 B).

**Table 1:** Ricker model parameter values for each phase between break points resulting from fitting population data of 1) multicolored Asian ladybeetles from Michigan, USA (1994-2017), and 2) the area occupied by monarch butterflies in their winter habitat in central Mexico (1995-2017). The parameter *r* is the per capita yearly intrinsic rate of increase and *K* is the carrying capacity (e.g., average number of adult ladybeetles captured per trap annually and hectares occupied by monarchs annually).

| Species | Years in subset | $r$ ( ± SE) | $K$ ( ± SE) |
| --- | --- | --- | --- |
| **Ladybeetle** | 1994-2000 | $1.3 \pm 0.3$ | $0.31 \pm 0.02$ |
| *Harmonia axyridis* | 2001-2005 | $2.3 \pm 0.3$ | $0.43 \pm 0.03$ |
| | 2006-2017 | $1.6 \pm 0.3$ | $0.27 \pm 0.03$ |
| **Monarch** | 1995-2003 | $1.0 \pm 0.5$ | $10.1 \pm 1.9$ |
| *Danaus plexippus* | 2004-2008 | $1.6 \pm 0.2$ | $5.6 \pm 0.3$ |
| | 2009-2017 | $1.2 \pm 0.4$ | $2.8 \pm 0.5$ |

**Figure 3:**
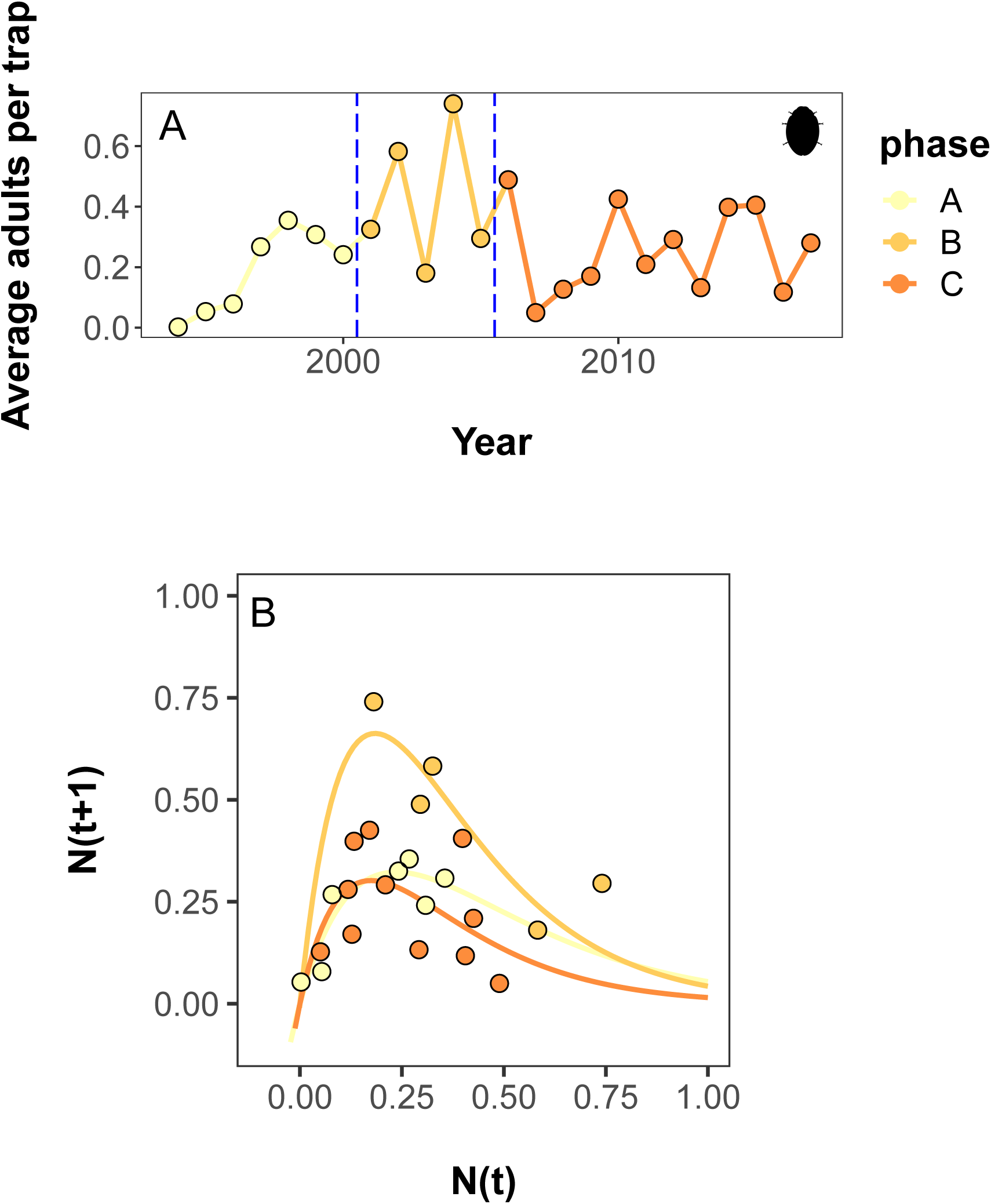
Dynamic Shift Detector breaks and Ricker model fits for an invasive species. Population data documenting the invasion of multicolored Asian ladybeetle in Michigan, USA from 1994-2017. A) Time series data showing the average number of adults captured, per trap, per year. Vertical blue lines indicate years in which dynamic shifts occurred, as estimated by the Dynamic Shift Detector algorithm. B) Ricker fits of time series data segments. Ladybeetle art by M. Broussard, used under a CC-BY 3.0 license.

Our results can be explained in the context of the known ecology of this ladybeetle. Dynamics of the ladybeetle invasion appear to be closely coupled with prey availability [41,43–45], which, in turn, is driven by documented pest management practices (neonicotinoid insecticide use; [42]) leading to a relatively simple pulsed change. The first shift in the dynamics of the Asian ladybeetle, after 2000, corresponds to the well documented arrival and establishment of soybean aphid to North America, a preferred prey item from the ladybeetle’s native range [46,47]. The invasion of this aphid dramatically increased resources available to the ladybeetle in habitats where the beetles were already well-established [40], supporting both a higher carrying capacity and a greater intrinsic growth rate. The second shift, after 2005, was less strongly supported, but coincides with the introduction and uptake of a management strategy for aphids that incompletely controlled the prey item. Landscape-scale use of neonicotinoid insecticides decreased prey numbers, particularly during the spring when aphids colonize new hosts, which could limit early season reproduction of ladybeetles [42]. Indeed, in this case, we would expect a weaker shift in dynamics as the prey item is incompletely controlled, and control tactics were not uniformly adopted across the prey’s range simultaneously.

#### Monarch butterflies in Mexican overwintering grounds

The eastern population of the North American monarch butterfly (*Danaus plexippus*) is migratory, with the majority of individuals overwintering in large aggregations in Oyamel fir forests within the transvolcanic mountains in the central region of Mexico [48]. Monarchs are highly dispersed over their breeding season, occupying landscapes throughout the agricultural belt in central and eastern United States and southern Canada [49]. As such, estimates of the overwintering population size can provide a convenient and inclusive annual metric of the size of the eastern migratory population [50]. This population of monarchs has been in dramatic decline in recent decades, although the degree and cause of this decline is hotly debated [51]. We used data on the total area occupied by monarchs from 1995-2017 (based on early winter surveys conducted in December) compiled by the World Wildlife Fund Mexico (available at MonarchWatch; Lovett 2017).

The DSD algorithm estimated that the best break point combination fit for the monarch overwintering data was a single break after 2003 (Fig. 4; AICc=120.18). However, the algorithm indicated that two additional break point combinations, a single break after 2006 (AICc=121.87) and a two-break combination of 2003 and 2008 (AICc=-121.86), had equivalent performance. The weight analysis computed weights of 0.49, 0.14, and 0.26, for 2003, 2006, and 2008 respectively, suggesting that the break at 2006 is erroneous and providing intermediate support for the 2003 and 2008 breaks. As with our ladybeetle case study, the strength of evidence was strongest for the first break, and weaker for the second break. The shift corresponds with a >50% reduction in K in 2003, and, if the secondary break is taken at 2008, a further reduction of K nearing 50% again at that point (Table 1; Fig. 4 B).

**Figure 4:**
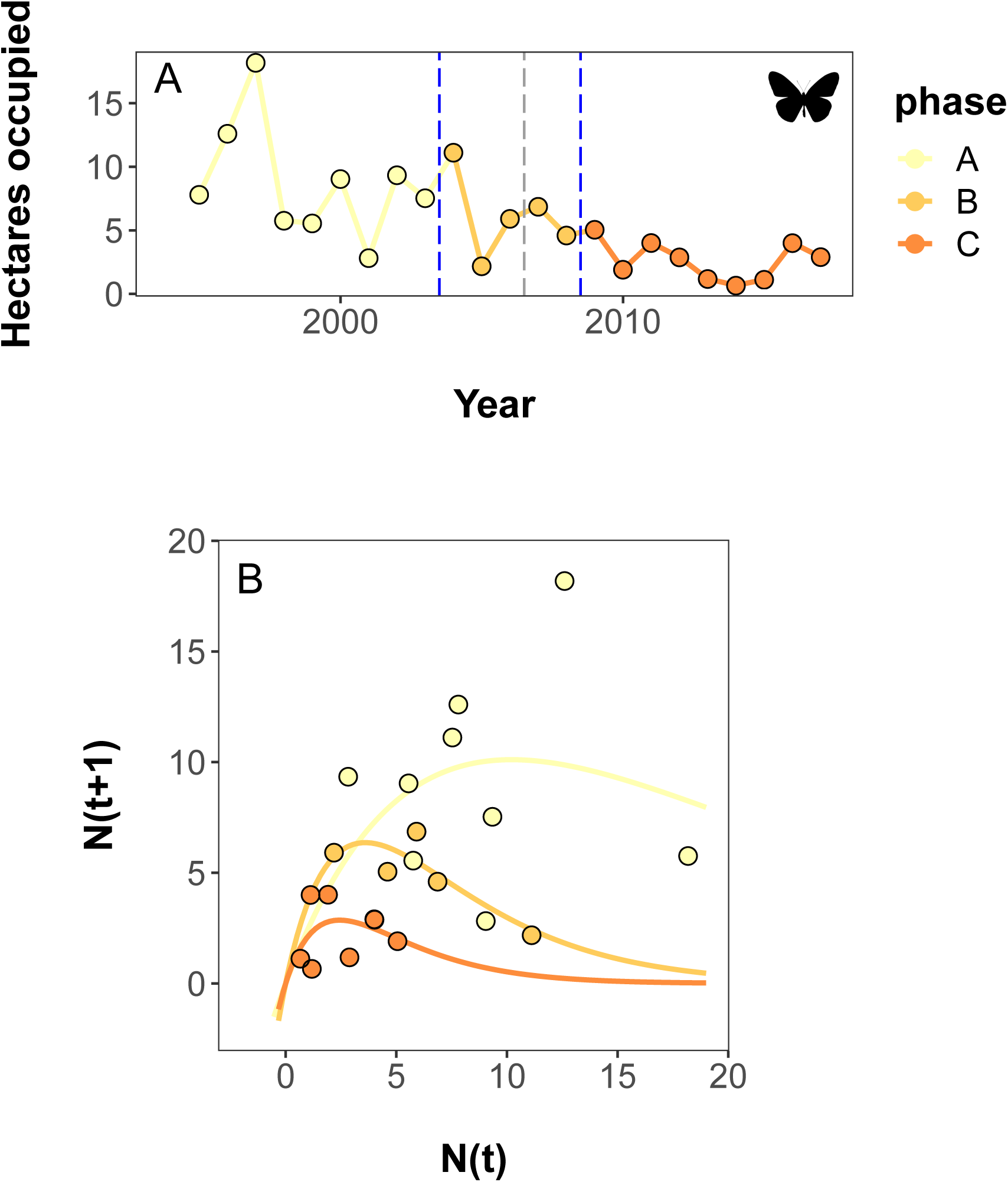
Dynamic Shift Detector breaks and Ricker model fits for a species of conservation concern. Population data documenting the area occupied by monarch butterflies in their winter habitat in central Mexico from 1995-2017. A) Time series data showing the total area occupied by overwintering monarchs each year in December. Vertical blue lines indicate years in which dynamic shifts occurred, as estimated by the Dynamic Shift Detector algorithm. B) Ricker fits of time series data segments. Butterfly art by D. Descouens and T.M. Seesey, used under a CC-BY 3.0 license.

The patterns we observe are consistent with a leading hypothesis to explain monarch population decline. Loss of milkweed hostplants due to changing agricultural practices on Midwestern breeding grounds [53,54] is hypothesized to be a major driver in the dynamics of this species. Changing herbicide practices in central North America have largely eliminated milkweed hostplants from agricultural field crops, with fairly consistent, low levels of milkweed on the landscape starting from about 2003-2005 [55]. Although glyphosate tolerant soybeans and maize were introduced to the US market in 1996 and 1998 respectively [56], actual glyphosate use lagged behind, with dramatic increases in use of the pesticide in 1998-2003 in soybean, and 2007-2008 in maize [57].

However, additional drivers likely play a role in monarch processes given the uncertainty in our results. Abiotic drivers of monarch population dynamics are complex and can interact at local, regional, and continental scales [58]. Other studies have implicated climate [59], extreme weather events [60], changing habitat availability on wintering grounds [61], and mortality during the fall migration [62,63] as possible factors influencing monarch population dynamics. With many super-imposed drivers, monarch dynamics are likely driven by both press and pulsed processes, making the detection of discrete break points associated with dynamic shifts complicated.

## Discussion

The DSD algorithm provides a novel tool for evaluating shifts in parameter values that govern density-dependent populations, such as carrying capacity and population growth rates. As illustrated with our simulations and case studies, the DSD algorithm can not only identify and quantify parameter changes but also assess uncertainty in potential break points and help detect time frames where additional research should be focused. We recommend that the model selection approach be used to identify a list of potential break points and break point combinations, and the weighting tool be used to evaluate the strength of evidence for each potential break, providing a clear direction to focus downstream research on changing dynamic processes. Characterizing dynamic transitions in population time series data has been hindered by a lack of a common, accessible, and empirical approaches [2]. Most assessments of break points in ecological research are *ad hoc* [12,14–17], introducing the potential for bias in break point selection, particularly in cases where nonlinear dynamics are likely to occur. Our DSD algorithm directly addresses this gap in tools, using an information-theoretic, model selection approach and a framework that can incorporate a variety nonlinear population models to assess ecological processes.

The core novelty of our tool lies in the model selection procedure used within the DSD algorithm, which allows for greater accuracy than common break point detection models [1]. The DSD additionally allows users to assess the confidence in a given break point, as well as providing a measure of which break points are likely to appear together, if multiple dynamic shifts have occurred. Information-theoretic approaches such as model selection using AIC, may be prone to over-fitting, particularly when data are limited [35]. Thus, we used AICc, the Akaike Information Criterion corrected for small sample sizes, as the selection criterion within the DSD algorithm. AICc is generally recommended in place of AIC in situations where small samples sizes are being examined as it incorporates a penalty term that is inversely related to the number of observations. As sample size increases, the penalty for model complexity is reduced and AICc approaches AIC [35]. Break point detection methods must incorporate a compromise between sensitivity and penalty for over-parameterization [1]. We examined the performance of both AICc (here) and AIC (in Appendix S1) and found that using AICc performed best for our simulated data. The DSD algorithm aides in the interpretation of break points by incorporating a metric based on Akaike weights, which allows users to assess the relative ‘strength’ of multiple breaks. Where many tools aim to identify points at which parameter changes occur, the DSD algorithm provides a measure of the confidence in each break, as well as a measure of how differing break sets perform in explaining variation in the data relative to each other.

The performance of the DSD algorithm was fairly stable among the break point simulations we tested. We found that the amount of environmental/sampling noise (ranging from 1-15% of the population size) had little effect on algorithm performance (Fig. 1 A). Other input conditions had relatively greater impacts on the performance of the DSD algorithm, depending on which parameter was changed and by how much. Large shifts in a population’s carry capacity were more easily detected than smaller shifts (Fig. 1 C). However, large changes in population growth rate were harder to detect, but this effect was most pronounced when simulated data contained multiple breaks (Figs. 1, 2D). Although larger shifts in regression parameters would, intuitively, lead to a higher likelihood of detection, large shifts in growth rate are also more likely to induce variations in transient dynamics in the years immediately following the shift, potentially making the timing of shifts more difficult to pinpoint. Similarly, longer time series yielded results that were more error prone (Fig. 1–2E). This is likely because there were simply more possible break-point combinations for the algorithm to select from and because the penalty for increasing parameterization (i.e. AICc) decreases as sample sizes grow (leading to increasing likelihood of identifying extra, erroneous breaks).

In our case studies, we found interpretation of the ladybeetle example was straightforward (Fig. 3). Our top break point combination and the equivalently-performing set did not contain contradictory information: each candidate set was simply a subset of breakpoints from the most complex set, and only two break points were found. Both of these break points were associated with moderate or greater weights, although the values of these break weights were in the intermediate range (i.e., between 0.2 and 0.8), suggesting breaks in natural systems may not be as well behaved as those in simulated data. The monarch butterfly case study results were slightly more ambiguous, as the model selection tool identified a break that the weighting tool suggested was erroneous (Fig. 4). Weights of the two most strongly-supported breaks were numerically similar to those of the ladybeetle case study, and are also interpretable with knowledge of the study system. However, the model selection results suggest additional, superimposed processes may be affecting monarch population dynamics, creating a noisier signal.

The DSD algorithm is readily adaptable to other population models and, indeed, potentially to other nonlinear processes. Density-dependent population growth has the potential to mask transition points because of its inherent nonlinear structure. For example, transient dynamics occurring immediately after a temporary disturbance can result in a change in population size, but not necessarily in the rules governing dynamics. We used the Ricker model as the core population model within the algorithm because it had a number of useful characteristics, namely its simple parameterization and realistic behavior [27]. However, simple density-dependent population models including Ricker, Beverton-Holt, and logistic models have similar performance in predicting outbreaks of insects driven by food limitation [64]. Indeed, in early iterations examining the DSD, we fit both logistic and Ricker models to real data and found that the two models ranked break point combinations nearly identically, even while the Ricker model generally provided a better fit for the data [42]. Thus, we expect the DSD algorithm would have similar performance across models with similar structures, but performance may vary with other model structures, particularly those that incorporate additional terms.

We recommend users carefully consider the strengths and limitations of the DSD algorithm in the context of their own data. For example, if changes to parameter values occur frequently (e.g., less than 3-4 years or time periods), the frequency of shifts would violate the constraints placed on the DSD to prevent overfitting. We also observed that the likelihood of identifying erroneous break points increased as time series length increased. Thus, in cases where a long time series exists, but a particular time period is of interest, the DSD algorithm could be used on the time period of interest alone to minimize the likelihood of distracting or erroneous results. The DSD algorithm functions as a method for identifying break points within time series data and quantifying the strength of evidence for each potential break point. When interpreted in the context of species biology, the DSD algorithm has the potential to aid management decisions, identify critical drivers of change in species’ dynamics, and help determine where best to focus additional research efforts.

## Supporting information

Supplemental Information 1- Analysis using AIC

Supplemental Information 2- Function Descriptions

## Acknowledgements

The conception of an earlier version of this algorithm came about out of conversations with Wopke van der Werf and Douglas Landis, and the algorithm has incorporated feedback from colleagues from the US-Long Term Ecological Research network throughout its development.

## Supporting Information

S1– Additional Simulations using AIC for break point selection

S2– Description of functions used in the Dynamic Shift Detector (DSD) algorithm

