## Supplemental Information 1- Analysis using AIC for "The Dynamic Shift Detector: an algorithm to identify changes in parameter values governing populations"

**S1- Additional Simulations using AIC for break point selection**

**Methods**

In this appendix, we present a detailed examination of the behavior of the Dynamic Shift Detector (DSD) algorithm using an alternate information criterion with less conservative behavior: AIC. In our initial calibrations of the DSD algorithm, we found that AIC was generally more sensitive to identifying break points, but also tended to over-fit (i.e. include extra breaks), likely because model complexity is not penalized as severely as with AICc. For comparison purposes, we conducted an identical simulation study to the one described in the main text with AICc. We also analyzed the case study data using AIC criterion and present the results herein.

The AICc-based DSD algorithm in the main text is a compromise between sensitivity and specificity. In general, we recommend users of the DSD algorithm use AICc with their data. However, there may be cases where it is desirable to gain a more liberal estimate of dynamic shift changes. In such case, the more sensitive AIC can be used to rank break point combinations. Results of the DSD algorithm using AIC and AICc were qualitatively similar, except that using AIC led to the inclusion of more possible breaks with higher break weights of both true and erroneous breaks. Thus, this more sensitive approach may be most useful in the context of hypothesis generation, rather than as an explicit hypothesis test.

*Simulation study*

We examined the dynamic shift detector’s performance for all test scenarios described in the main text using the AIC criterion. First, we evaluated the ability of the algorithm to detect scenario initialization conditions within the set of equivalent break point combinations (Fig S1-1). We also examined the performance of the break-point weighting tool from the perspective of its average weightings of true and erroneous break points (Fig. S1-2).

In general, the AIC-based model sets were less likely to identify true breaks within the equivalently performing break point combination sets in comparison to using AICc. The performance of the most complex parameterizations were similar between AIC and AICc (accuracy of about 70%), but the performance associated with all other parameterizations was greatly reduced with AIC (<60%; Fig. S1-1). We observed the same general patterns around the DSD’s performance in parameter space: decreased performance with increasing experimental noise, extreme initial values of r, small changes in K, large changes in r, and longer time series length (Fig. S1-1).

With the weight analysis, we found that (as with AICc) the average weight of a true break (i.e. one that was intentionally simulated in the data) typically exceeded a value of 0.8 in the vast majority of parameterization cases (Fig. S1- 2). However, unlike the AICc results, erroneous breaks had higher average weights, with values up to approximately 0.4.

*Case study: ladybeetles*

As with the DSD analysis using AICc, the AIC analysis found two break points, one occurring after 2000 and one occurring after 2005, in the top break point combination model. However, the AIC analysis did not find any additional break point combinations with equivalent performance. Break weight analysis suggested a weight of 0.95 for the 2000 break, and a weight of 0.64 for the break after 2005. In the context of the AIC analysis, we would thus conclude unambiguous support for the break occurring at 2000 and moderately high support for the break at 2005, similar to results from the AICc analysis.

*Case study- monarchs*

For the monarch butterflies, using AIC as the information criterion in the DSD resulted in a much more complex result as compared to the AICc based analysis. In this analysis, a break point combination with two breaks (at 2003 and 2008) was selected as the top model (AIC=106.86), but there were four additional break point combinations with equivalent performance (1998, 2003, 2008, AIC=106.90; 1999, 2003, 2008, AIC=108.02; 2003, 2007, AIC=108.51; and 1998, 2003, 2007, AIC=108.54). Thus, the model suggested many more candidate break points (1998, 1999, 2003, 2007, and 2008) but the weighting analysis found variable strength of evidence for these breaks (0.41, 0.23, 0.81, 0.27 and 0.65, respectively). We conclude there is weak evidence for breaks at 1998, 1999, and 2007, moderately high evidence for a break at 2008, and fairly strong evidence for a break at 2003, which parallels results in the main text using AICc.

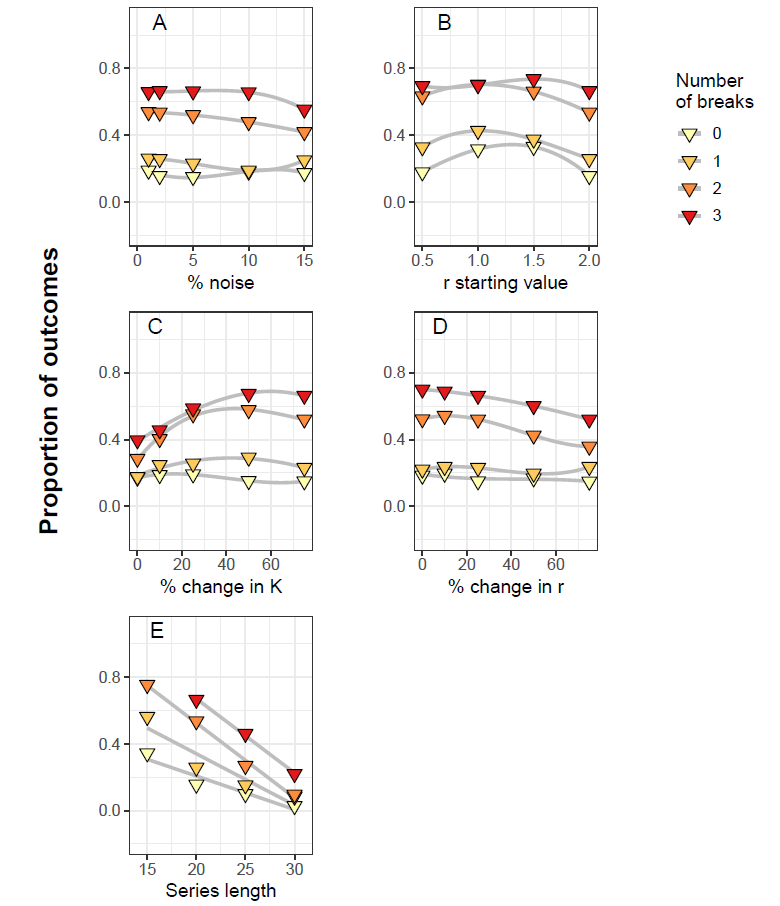

Figure S1-1: **Performance the dynamic shift detector model under varying conditions using AIC criterion.** Proportion of simulation results in which the true breaks were detected within the top break point combinations as identified by the DSD model implemented with an underlying Ricker model with varied A) noise (in the form of normally distributed error), B) starting values of the *r* parameter, C) percent changes in the *K* parameter, D) percent changes in *r*, and E) simulated time series length. Sets of 0, 1, 2 and 3 break points were randomly generated from within the set of possible values, and 250 datasets were simulated for each scenario. In each panel, other variables (that were not being varied) were held constant at their base values (i.e., noise=2%; starting value of *r* = 2; change in *r* = ±25%; change in *K* = ±75%; time series length = 20 years). Trends within a set of scenarios (grey lines) are illustrated with a third-order GAM smoothing line.

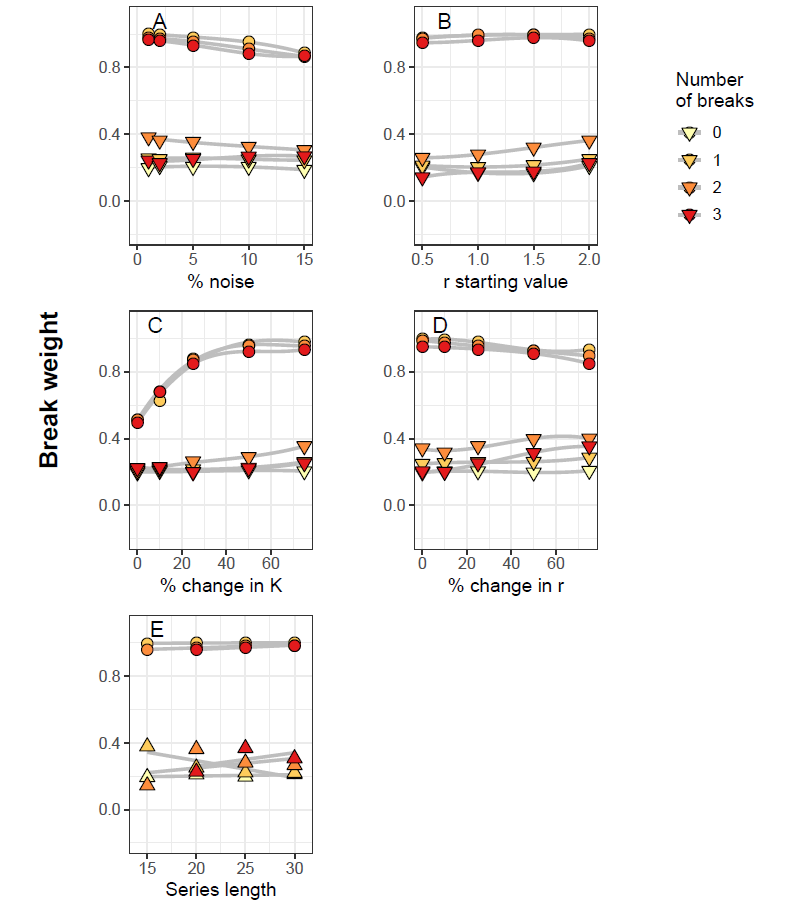

Figure S1-2: **Average break weight of break points found under varying parameterization conditions using AIC criterion.** Average weights of break points identified by the Dynamic Shift Detector model reflecting true parameterization conditions (diamonds) or erroneous breaks suggested by the model (triangles) under varied A) noise (in the form of normally distributed error), B) starting values of the *r* parameter, C) percent changes in the *K* parameter, D) percent changes in *r*, and E) simulated time series length. Sets of 0, 1, 2 and 3 break points were randomly generated from within the set of possible values, and 250 datasets were simulated for each scenario. In each panel, other variables (that were not being varied) were held constant at their base values (i.e., noise=2%; starting value of *r* = 2; change in *r* = ±25%; change in *K* = ±75%; time series length = 20 years). Trends within a set of scenarios (grey lines) are illustrated with a third-order GAM smoothing line.
