## Supplemental Information 2- Function Descriptions for "The Dynamic Shift Detector: an algorithm to identify changes in parameter values governing populations"

**S2- Description of functions used in the Dynamic Shift Detector (DSD) algorithm**

addNt1 - takes a raw data frame with columns for year, population abundance measure, and converts it to a three-column data frame with year, population in year, population in the next year, and gives the data consistent column names for use in downstream functions. Most of the functions described below require this function to transform data prior to use, unless otherwise specified.

AICcorrection - takes a data series and the number of breaks used in a given fit to calculate the AICc correction factor to be added to the total AIC for the fit.

Rickerfit - fits the Ricker model using the Levenberg-Marquart nonlinear least squares method for nonlinear model fitting to a given data frame. This function calls the nlsLM function from minpack.lm (Elzhov, Mullen, Spiess, & Bolker, 2016). To aide in model convergence, the function also computes a realistic starting value for K by calculating the mean of the population (based on the assumption that a population being fit to the Ricker model is likely fluctuating around its carrying capacity). The starting value for r was set at 1.5. The function outputs a vector containing the AIC, the estimate for r, its standard error, and the estimate for K and its standard error.

splitnfit - takes a given data fame, fits the complete data series with the rickerfit function, then subsets it in two by creating a break point three years after the start of the series, calls the rickerfit function to fit the data from each subset produced there. Then the function walks through the data, increasing the break point by one time step each iteration, and compiles the AICs and break points used for each fit, resulting in a data frame of break point combinations and respective AICs.

findbreakable - examines the output from the splitnfit function to determine if any of the break point combinations produced might be further subdivided (i.e., has enough points to not violate the rule we set to only fit series with four or greater points).

subsequentsplit - uses output from findbreakable function to identify cases where data can further be subsetted using the splitnfit function, feeds those cases in, and compiles results together with that produced by simpler break point combinations produced by splitnfit.

nbreaker - uses splitnfit, findbreakable, and subsequentsplit, combined with input data, to create a data frame consisting of a column of all possible break point combinations, and the respective AICs of the resultant fits. This function uses an iterative approach to allow simpler functions that break a data into two parts to be used to find an unlimited number of break points (within constraints of series length).

AICtally - takes data in, subjects it to nbreaker, pulls out the AICs produced by nbreaker, adds them together and counts the number of fits performed, number of breaks in the data, computes the corrected AICc using AICcorrection, and returns these values as a data frame.

allfits - appends the results of nbreaker and AICtally together into a single data frame, resulting in summary statistics for all possible break point combination fits for the input data series.

equivalentfit - takes in data, feeds it to allfits, and uses the output from allfits to pull out the subset of all equivalently-performing breakpoint combination fits (here, within 2 units of AICc or AIC, depending on user input), and outputs these fits as a data frame.

bestfit - feeds data to the equivalentfit function to get a data frame describing equivalent fits, and selects the one with the lowest AICc/AIC to output the specifics of that break point combination as a data frame.

bestmodel - feeds data to the bestfit function to identify the best break point combination, and then use that information to create a data frame describing the parameter estimates (r, K and standard error for each) for fitting the Ricker model to each of the subsets of timeseries, allowing a user to quantify the dynamic rule changes found by fitting the model changes at each break point.

modelspecification - this function takes data in the format produced by the bestfit function and produces a data frame describing the model specification of the given break point combination, for the case when a user wishes to investigate specification of similarly ranked models.

DSdetector - uses the raw time series data to produce a report, calling all the previous functions, either directly, or through other functions, with short explanatory text preceding each result. First, a simple plot of population over time is produced (N(t) by t), then data is fed to the addNt1 function, and the resultant N(t), N(t+1) data is plotted to visualize the potential for the data to conform to a Ricker curve. The data is fed through the allfits function, producing a complete list of all break point combinations tested and their respective fit statistics. The data is subsequently fed through the equivalentfit and bestfit functions so that a user can assess how the decision rules specified impacted the selection of the best model. Finally, the data is fed through the bestmodel function to produce the set of regression parameters for each time series subset produced by the best break point combination found.

allweights - uses allfits to compute Akaike weights of all break point combinations tested, based on AIC/AICc, depending on user input. Culls out break point combinations with Akaike weights of less than 0.001 to save processing time. Outputs a data frame of break point combinations with their respective Akaike weights.

breakweights - uses allweights to compute a relative weight for all individual breaks found, based on (Burnham & Anderson, 2002)Relative Variable Importance. For each prospective break, sums the Akaike weights for each break point combination this break appears in, compiles a data frame of break, weight. Because of approximation used in allweights, weights are normalized by dividing by the computed weight of the break at the end of the series (by definition, should be 1) Outputs a data frame of break, the computed weight and the corrected weight for each break occurring in models with Akaike weights >0.001.
